# Mathematical model for force and energy of virion-cell interactions during full engulfment in HIV: Impact of virion maturation and host cell morphology

**DOI:** 10.1101/859355

**Authors:** Elizabeth Kruse, Tamer Abdalrahman, Philippe Selhorst, Thomas Franz

## Abstract

Viral endocytosis involves elastic cell deformation, driven by chemical adhesion energy, and depends on physical interactions between virion and cell membrane. These interactions are not easy to quantify experimentally. Hence, this study aimed to develop a mathematical model of the interactions of HIV particles with host cells and explore the effects of mechanical and morphological parameters during full virion engulfment. The invagination force and engulfment energy were described as viscoelastic and linear-elastic functions of radius and elastic modulus of virion and cell, ligand-receptor energy density and engulfment depth. The influence of changes in the virion-cell contact geometry representing different immune cells and ultrastructural membrane features and the decrease in virion radius and shedding of gp120 proteins during maturation on invagination force and engulfment energy was investigated. A low invagination force and high ligand-receptor energy are associated with high virion entry ability. The required invagination force was the same for immune cells of different sizes but lower for a local convex geometry of the cell membrane at the virion length scale. This suggests that localized membrane features of immune cells play a role in viral entry ability. The available engulfment energy decreased during virion maturation, indicating the involvement of additional biological or biochemical changes in viral entry. The developed mathematical model offers potential for the mechanobiological assessment of the invagination of enveloped viruses towards improving the prevention and treatment of viral infections.

## 1. Introduction

For the human immunodeficiency virus (HIV) to establish the infection of a host cell, it employs a complex series of actions, all while evading any immune response in the host (Wilen et al. 2012). The first steps in this infection and replication cycle are receptor binding, membrane fusion, and entry into the host cell. The mechanisms of these initial steps have a critical impact on the entire infection process of HIV and could guide the formulation of treatment therapies (Gorai et al. 2021; Wilen et al. 2012). Currently, a major target in the search of viral therapeutics is the inhibition of the spike protein, which binds to the virus to the receptor of the host cell (Choudhary et al. 2021).

While the mechanism of HIV entry into host cells has been extensively studied from a biological and biochemical perspective (Miyauchi et al. 2009; de la Vega et al. 2011; Zaitseva et al. 2017; Coomer et al. 2020), the biophysical factors that may affect the mechanics of virus-cell interaction remain unclear (Sun and Wirtz 2006).

For example, mature HIV particles are known to have a higher entry ability into host cells compared to immature virions (Murakami et al. 2004; Wyma et al. 2004). Pang et al. (2013) reported that this increased entry ability of the virions into the cells, was the result of a reduced virion stiffness which was previously shown by Kol et al. (2007) to decrease upon virion maturation.

Biomechanical factors, such as the stiffness of the plasma membrane, cytoskeleton and cytosol of the target cell, can also influence the virus-cell mechanical interaction in addition to biochemical factors. Early viral endocytosis is driven by chemical adhesion energy that facilitates elastic deformation of the cell, which depends on the mechanical properties of the cytoplasm and cell membrane (Gefen 2010). Considering the physical interactions between virion and cell membrane, contact mechanics likely plays a large role in virion entry into the host cell.

Mathematical modelling of the mechanical interactions between virion and cell membrane allows for quantifying forces and energies involved in virion engulfment that are not easily accessible with experimental approaches. Such mathematical models have been used to investigate how contact force, mechanical work, and pressure varied with the engulfment depth of the virion (Gefen 2010; Sun and Wirtz 2006). Gefen (2010) studied the impact of virion size and cell stiffness on the forces, work and pressures during virion engulfment, limited to small cellular deformations. Both studies assumed a uniform global radius of the host cell (which is very large compared to the radius of the virion) and did not account for morphological irregularities on the cellular surface at the micro- or nanoscale. However, electron microscopy images of the HIV-cellular interaction (Gentile et al. 1994) reveal a cellular surface with localized curvatures in the nanometre range. Such localized surface morphology of the cell membrane may indeed play a role in the virion-cell interactions during engulfment and endocytosis and the likelihood of viral infection of a cell.

There is limited data on the role of mechanical forces in virus engulfment and the interplay between engulfment mechanics and the morphology and stiffness of the virion and the host cell. The current study aims to develop a mathematical model that allows investigating the sensitivity of the mechanics of virion engulfment to changes in morphological and mechanical characteristics of HIV virions and host cells towards guiding future experimental studies.

## 2. Methods

The virion engulfment model describes the engulfment energy and the invagination force and was implemented in MATLAB R2014a (MathWorks Inc, Natick, MA, USA). The model was based on continuum models for receptor-mediated endocytosis of viruses that employ contact mechanics and consider ligand-receptor complex formation energy (Sun and Wirtz 2006; Gefen 2010). All images in the Results section were produced using the export_fig toolbox for MATLAB (Altman and Wooford 2014).

The model assumes a virion-cell arrangement illustrated in Figure 1, with a virion radius very small compared to the cell radius, i.e. *R*_v_ << *R*_c_. Due to the large size difference between the cell and virion, the cell membrane was approximated as a flat surface in the region of the virus-cell contact. The infection process is initiated when the virion ligands (i.e. gp120) dock to the cell receptors (i.e. CD4), and the cell starts to engulf the virion to an engulfment depth, *d*, by generating an invagination force, *F*. The model also assumes an equal density of uniformly distributed and immobile ligands and receptors and does not include receptor diffusion considered in kinetic models such as Yi and Gao (2017). Further, the fluidity of the cell membrane was not considered, and the cell membrane represented as a continuum element with membrane tension. The engulfment process is a critical initial step in triggering fusion between the virion and cell membrane and infecting the host cell.

**Figure 1.**
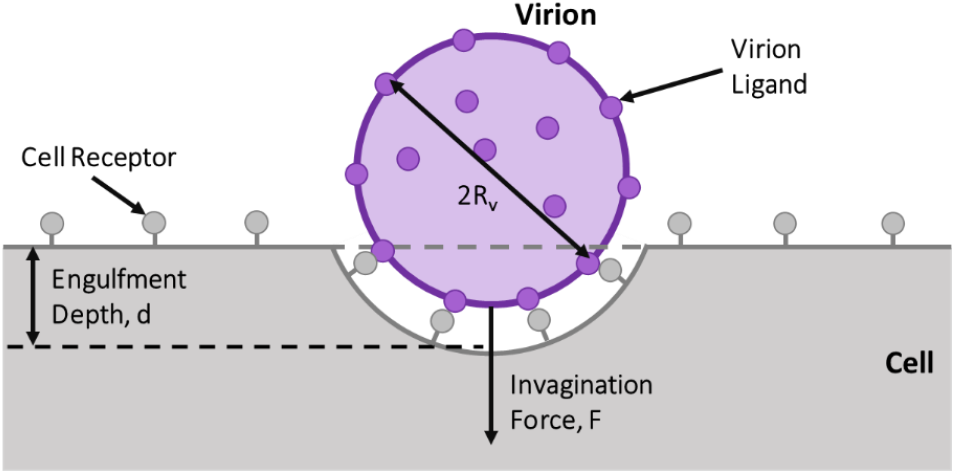
Schematic of virion-cell membrane interaction during virion engulfment.

### 2.1. Engulfment Energy

The total energy, *W*_T_ of the engulfment process comprised of three energy terms, namely the adhesion energy, *W*_1_, the membrane bending energy during the engulfment process, *W*_2_, and the energy of the cytoskeleton deformation, *W*_3_:

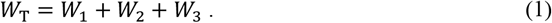

In Eq. (1), the adhesion energy, W_1_, is the energy gained from ligand-receptor binding that drives engulfment and is expressed as a negative value. The other two energy terms, i.e. the membrane bending and tension energy, W_2_, and cytoskeletal deformation energy, W_3_, are energies required during the engulfment process and are expressed as positive values. Thus, high entry ability is associated with a large negative value of the adhesion energy, representing a high magnitude of energy gained from ligand-receptor binding, and low energies required for membrane bending and cytoskeletal deformation.

#### 2.1.1. Adhesion Energy

The adhesion energy, *W*_1_, released by ligand-receptor binding, Eq. (2), is utilized to deform the cell membrane, thereby driving the entry process:

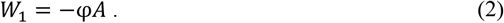

Here, φ is the ligand-receptor energy density function, and *A* is the contact area between the virion and the cell. This contact area, *A*, formed by a sphere (virion) and a half-space (cell) is a function of the engulfment depth, *d*, and the virion radius, *R*_*v*_, (Johnson 1987):

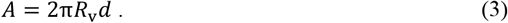

The radius of a mature and immature HIV virion is taken as *R*_v_ = 55 nm and 73 nm, respectively (Gentile et al. 1994).

The receptor-ligand density function, φ, depends on the energy gained per receptor-ligand complex, *f*, and the receptor complex density, ρ:

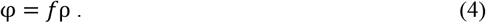

The energy gained, *f*, is 20*k*_B_*T* (Sun and Wirtz 2006; Li et al. 2018; Leckband and Israelachvili 2001), where *k*_B_ is the Boltzmann constant (1.3807 x 10^−23^ J/K), and *T* is the absolute temperature of 310 K for the human body (Gefen 2010). The receptor complex density, ρ, is determined by the ratio of the total number of glycoproteins and the virion surface area 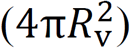. For HIV, each gp120 spike consists of 3 glycoproteins. Therefore, an immature HIV virion with 72 spikes (Gelderblom 1991; Grief et al. 1989) has 216 glycoproteins, and a mature HIV virion with 10 spikes (Layne et al. 1992) has 30 glycoproteins.

#### 2.1.2. Membrane Energy

The membrane bending and tension energy, *W*_2_, was assumed to be purely elastic energy and did not account for the viscoelasticity of the cytoskeleton and the heterogeneity of the cell. This energy is well described by the Canham-Helfrich theory (Canham 1970; Helfrich 1973) and is defined as:

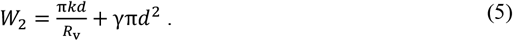

Here, *k* is the bending modulus of the cell membrane, with *k* = 20*k*_B_*T* (Wiegand et al. 2020; Deserno and Bickel 2003), and γ is the cellular surface tension with γ ≈ 0.005*k*_B_*T*/nm^2^ (Sun and Wirtz 2006).

#### 2.1.3. Cytoskeletal Deformation Energy

Two formulations for the energy of the cytoskeleton, *W*_3,_ are presented in this paper. The first was calculated using a linear-elastic function for small deformations, and the second as a viscoelastic function for the full engulfment process.

##### Linear-elastic Formulation

The elastic energy of the cytoskeleton, *W*_3el_, is a function of the effective elastic modulus, *E\**, the effective radius, *R\**, of the cell and the virus, and the engulfment depth, *d*:

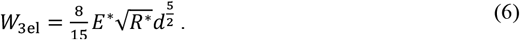

The effective elastic modulus is

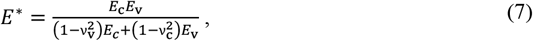

with the elastic modulus of the cell of *E*_c_ = 0.62 kPa (Fregin et al. 2019), and the elastic modulus of the HIV virion of *E*_v_ = 440 MPa for mature particles and *E*_v_ = 930 MPa for immature particles (Kol et al. 2007). The Poisson’s ratio of the HIV particle and the cell are ν_v_ = 0.4 (Ahadi et al. 2013) and ν_c_ = 0.5 (Sun and Wirtz 2006).

The effective radius, *R\**, is a function of the virion radius, *R*_v_, and the cell radius, *R*_c_:

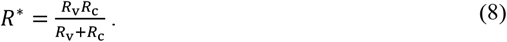

The radius of the entire cell and localized cell membrane morphology were used to represent cell dimensions, i.e. *R*_c_ = 4,000 nm for lymphocytes and *R*_c_ = 40,000 nm for macrophages (Kierszenbaum and Tres 2015), and *R*_c_ = 10 and 55 nm approximated from Gentile et al. (1994, Fig. 2).

**Figure 2.**
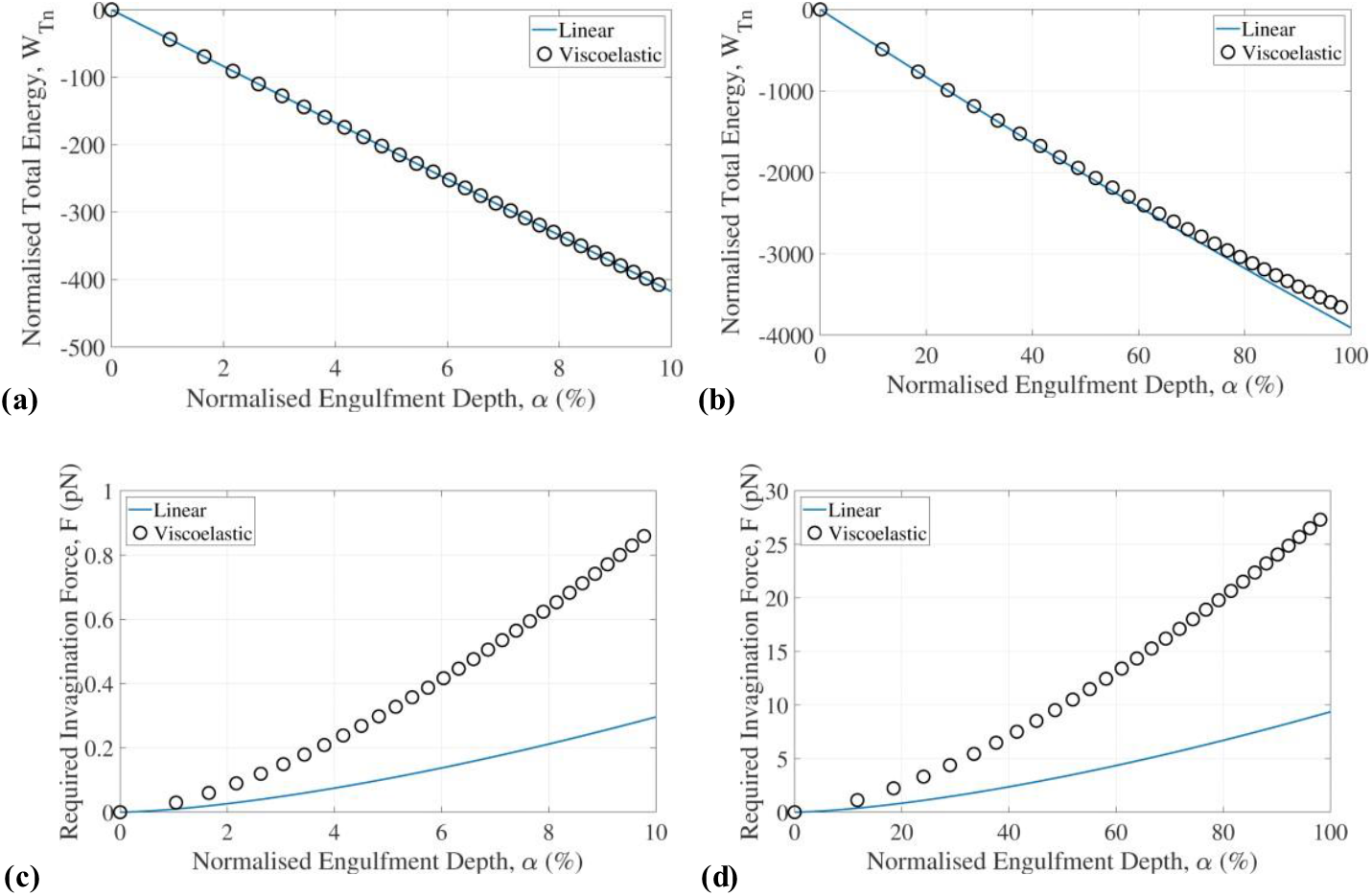
**The results of the viscoelastic model and linear-elastic model showing the total energy for (a) early (α ≤ 10%) and (b) full engulfment and the invagination force for (c) early (α ≤ 10%) and (d) full engulfment. For these results, the following model parameters were used: *R***_**c**_ **= 4 000 nm, *R***_**v**_ **= 55 nm, *E***_**v**_ **= 440 MPa, and ρ = 216/(4π*R***_**v**_)

##### Viscoelastic Formulation

To determine the viscoelastic energy *W*_3vis_, the Kelvin-Voigt viscoelastic model was used (Li et al. 2018; Micoulet et al. 2005):

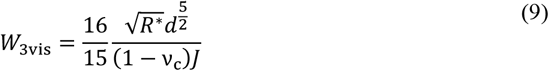

where *J* is a function of time, *t*, the elastic modulus of the cell, *E*_c_, and a function τ:

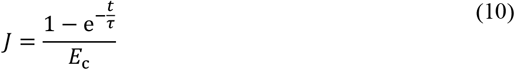

The function τ is the ratio between the viscosity, η, and elastic modulus, *E*_c_, of the cell:

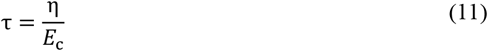

with a viscosity of the cell of η = 8.7 Pa (Fregin et al. 2019).

Assuming δ*W/*δ*d* = 0, the equilibrium engulfment depth function (Li et al. 2018) is:

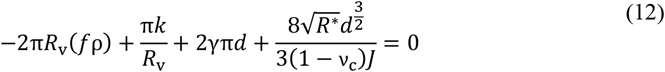

allowing to determine the relationship between engulfment depth and time.

### 2.2. Invagination Force

The invagination force can be calculated using either the assumption of linear elasticity for small deformations during early-stage engulfment or a viscoelastic function for the full engulfment of the virion. The force value indicates the force required to deform the cell around the virion, and thus a high force value suggests a lower entry ability.

#### 2.2.1. Elastic Force

The elastic force required for the invagination of the virus into the cell was defined using the Hertz model of frictionless contact between two spheres and can be derived from the elastic energy equation Eq. (6):

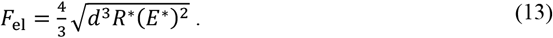

According to Dintwa et al. (2007), the assumption of frictionless contact is reasonable.

#### 2.2.2. Viscoelastic Force

The viscoelastic invagination force was determined using the Hertz contact model for a rigid sphere and a viscoelastic half-space. The equation can be derived from the viscoelastic energy equation found in Eq. (9):

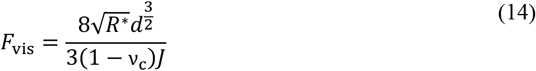

### 2.3. Data Normalisation

A normalized engulfment depth, α, is used to relate the engulfment depth of the virus to the virus radius:

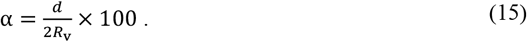

As the assumption of linear elasticity is only reasonable for small deformations, predictions with the linear-elastic model were limited to a normalized engulfment depth of α ≤ 10% (Gefen 2010).

The normalized total engulfment energy is obtained as:

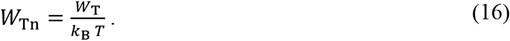

### 2.4. Parameter summary

Table 1 summarises the key parameters and their values used in the current study and those used in other studies. The first seven parameters listed have not been varied, whereas the last four parameters have been varied for sensitivity studies.

**Table 1.**
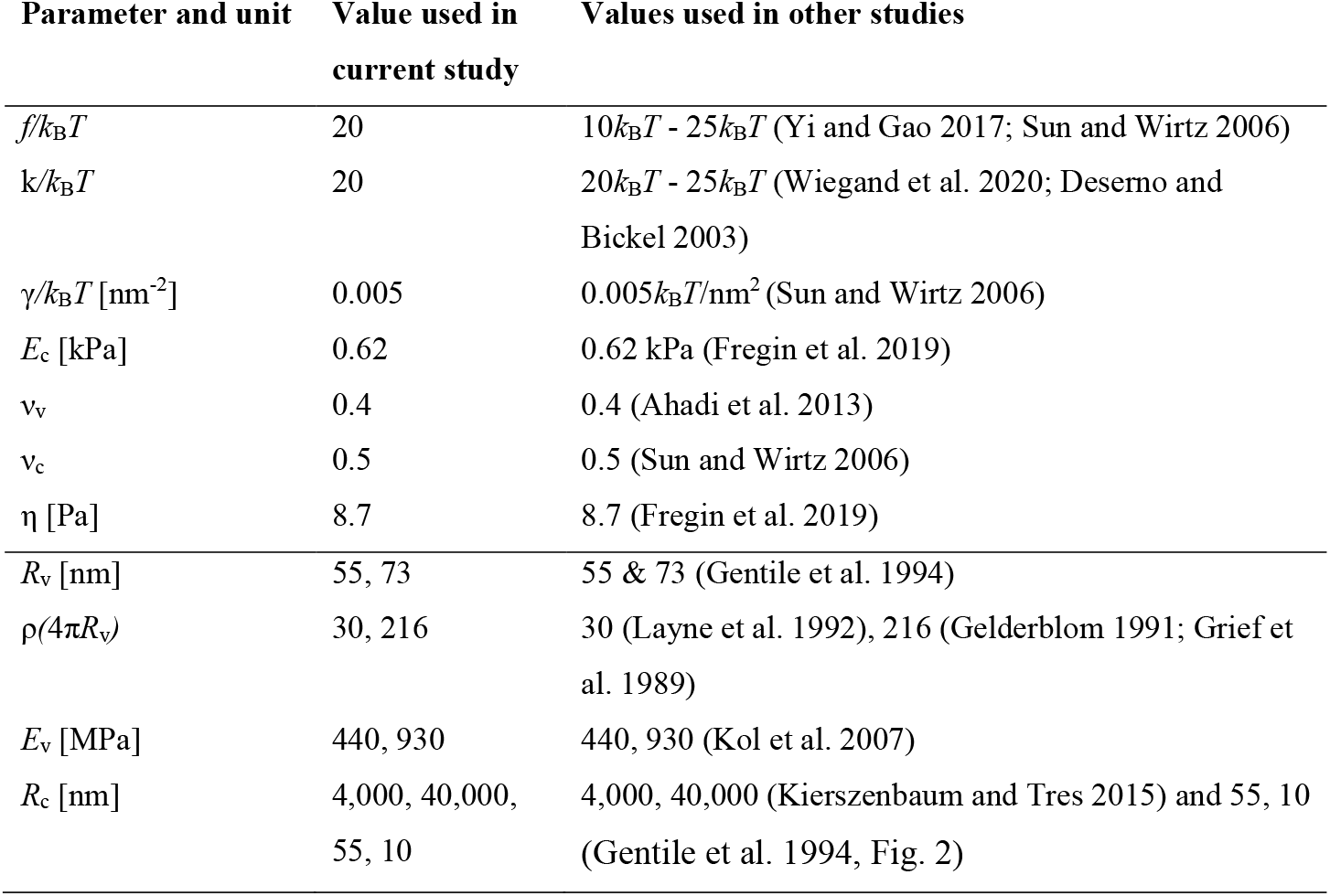
Model parameters and values used in this study and values from literature.

## 3. Results

### 3.1. Comparison of Linear-elastic and Viscoelastic Model Predictions of Engulfment Energy and Invagination Force

For the total energy, the linear-elastic model and the viscoelastic model yields similar results for early-stage engulfment with α ≤ 10%, whereas the linear-elastic model slightly underestimates the total energy for an engulfment with 10% < α ≤ 100% (Figure 2). For the invagination force, the linear-elastic model predicts substantially smaller values than the viscoelastic model at all stages of engulfment. Further comparative results for the linear-elastic and viscoelastic model will be reported in the following sections.

### 3.2. Cell Type and Morphologies

There is no discernible difference in the force required for the invagination of lymphocytes and macrophages with *R*_c_ = 4,000 and 40,000 nm, respectively, which are very large compared to a mature HIV particle with *R*_v_ = 55 nm (Figure 3). However, a substantial decrease in the required invagination force is observed when virion engulfment is considered in the region of localized cell membrane morphologies with a local cell radius (*R*_c_ = 10 and 55 nm) of the same order of magnitude as the virion radius (*R*_c_ ≈ *R*_v_). The reduction in the invagination force for *R*_c_ ≈ *R*_v_ is observed in the linear-elastic and the viscoelastic model. For 100% engulfment, the required invagination force was 28.3 pN for a macrophage with *R*_c_ = 40,000 nm and 28.1 pN for a lymphocyte with *R*_c_ = 4,000 nm, whereas it was 20.0 pN and 11.1 pN for local curvatures of *R*_c_ = 55 nm and 10 nm, respectively (Figure 3 c).

**Figure 3.**
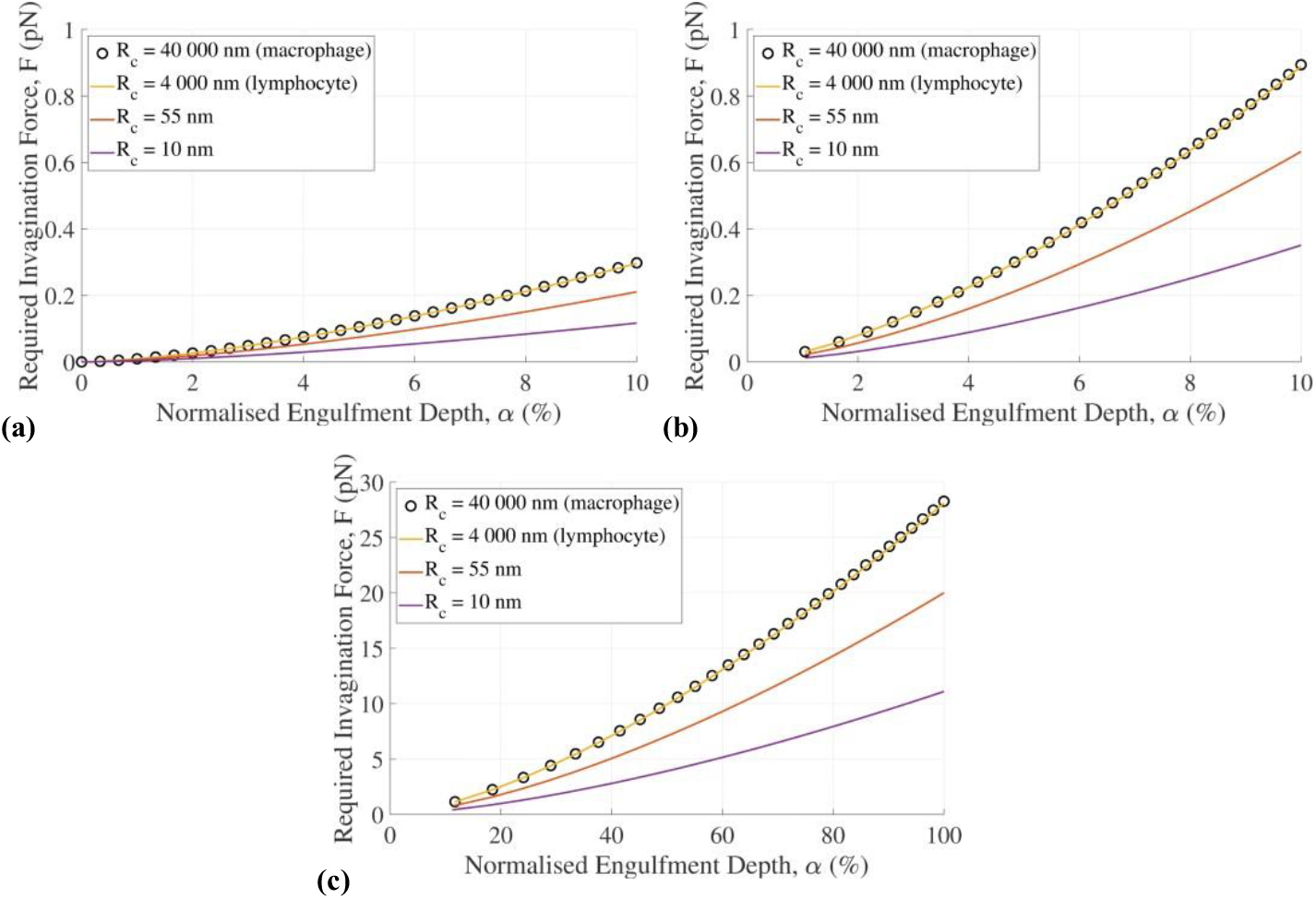
**Required invagination force versus normalized engulfment depth for different values of the cell radius representing a macrophage (*R***_**c**_ **= 40,000 nm), lymphocyte (*R***_**c**_ **= 4,000 nm), and localized cell membrane curvatures (*R***_**c**_ **= 10 and 55 nm). Results from (a) the linear-elastic and (b) the viscoelastic model for early stage engulfment (α ≤ 10%) and from (c) the viscoelastic model for full engulfment. The following virion parameters were used: *R***_**v**_ **= 55 nm, *E***_**v**_ **= 440 MPa, and ρ = 216/(4π*R***_**v**_)

The engulfment duration increases with increasing radius of the cell membrane. Full engulfment is predicted to complete in 2.97 ms for the macrophage (*R*_c_ = 40,000 nm), 2.95 ms for the lymphocyte (*R*_c_ = 4,000 nm), and 2.03 ms and 1.09 ms for the localized cell membrane curvatures of *R*_c_ = 55 nm and 10 nm (Figure 4). The engulfment speed decreases during the engulfment.

**Figure 4.**
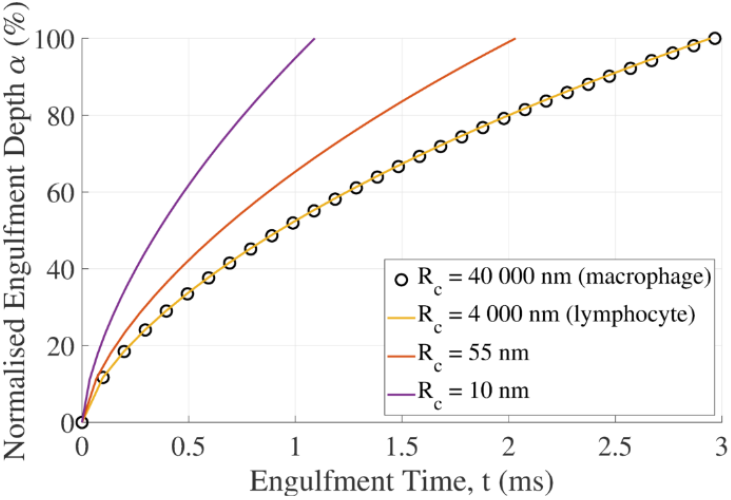
**Normalized engulfment depth of the virion versus engulfment time predicted with the viscoelastic model for different cell membrane radii representing macrophage (*R***_**c**_ **= 40,000 nm), lymphocyte (*R***_**c**_ **= 4,000 nm) and localized membrane curvatures (*R***_**c**_ **= 10 and 55 nm). Virion parameters used: *R***_**v**_ **= 55 nm, *E***_**v**_ **= 440 MPa, and ρ = 216/(4π*R***_**v**_)

### 3.3. Variation of Virion Size and Number of GP120 Spikes During Maturation

For the decrease in HIV virion size during maturation from *R*_v_ = 73 nm to *R*_v_ = 55 nm, there is no discernible difference in total engulfment energy during early-stage engulfment with α ≤ 10%, both for the linear-elastic and the viscoelastic model (Figure 5 Normalized total engulfment energy versus normalized engulfment depth for HIV particles with different radius and number of gp120 spikes related to virus maturation predicted with (a) the linear-elastic and (b) the viscoelastic model for early-stage engulfment, and with (c) the viscoelastic model for full engulfment. A cell radius of *Rc* = 4,000 nm was used

**Figure 5.**
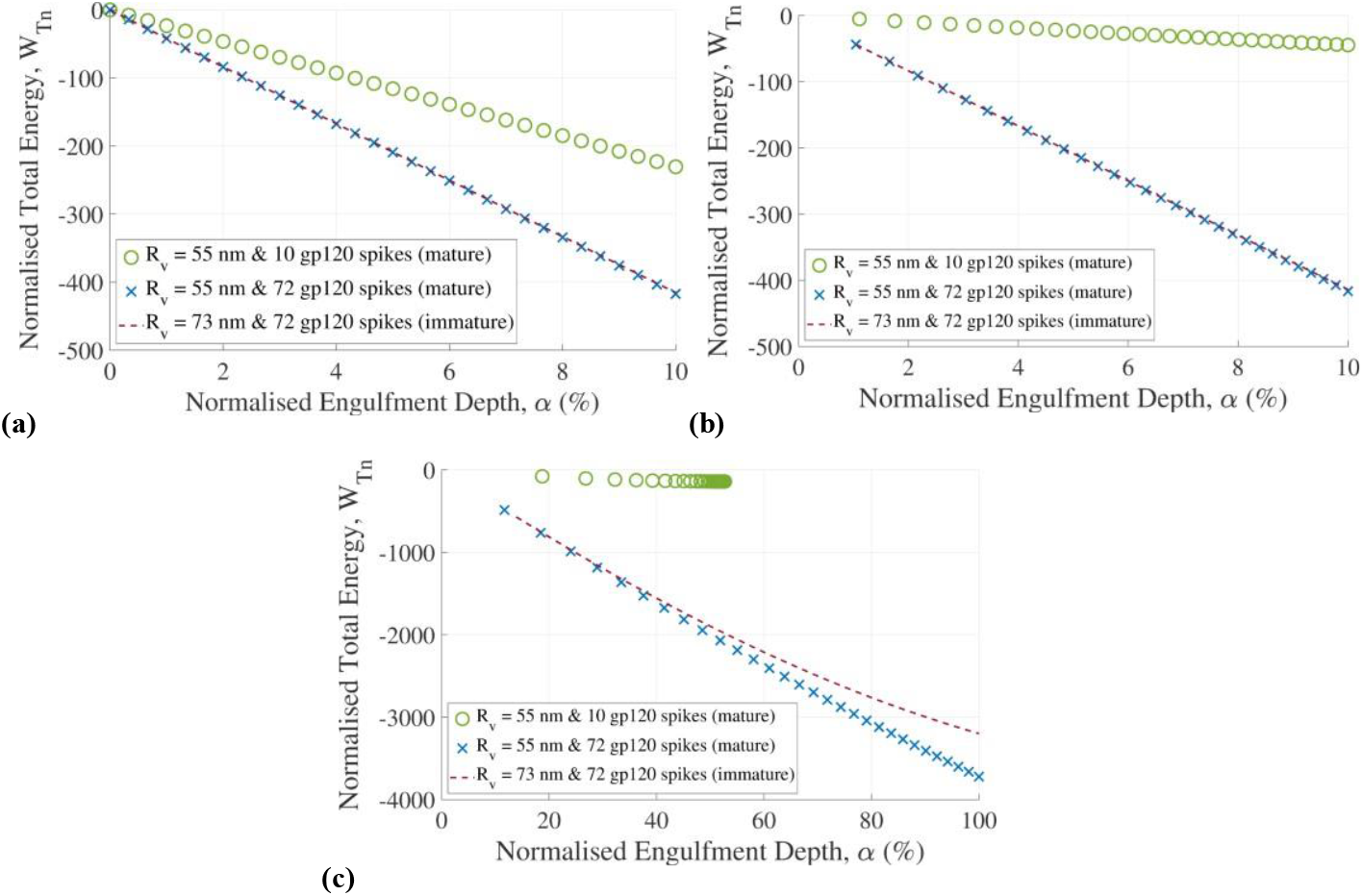
**Normalized total engulfment energy versus normalized engulfment depth for HIV particles with different radius and number of gp120 spikes related to virus maturation predicted with (a) the linear-elastic and (b) the viscoelastic model for early-stage engulfment, and with (c) the viscoelastic model for full engulfment. A cell radius of *R***_**c**_ **= 4,000 nm was used**

The duration of complete engulfment (α = 100%) is 8.88 ms for the immature virion and 2.88 ms for a mature virion with a smaller size (*Rv* = 55 nm versus *Rv* = 73 nm) but the same number of 72 gp120 spikes as the immature virion (Figure 6). Taking further into account the reduction in the number of gp120 spike proteins (10 versus 72) during maturation, the maximum engulfment is limited to 53% and reached after 64.62 ms.

**Figure 6.**
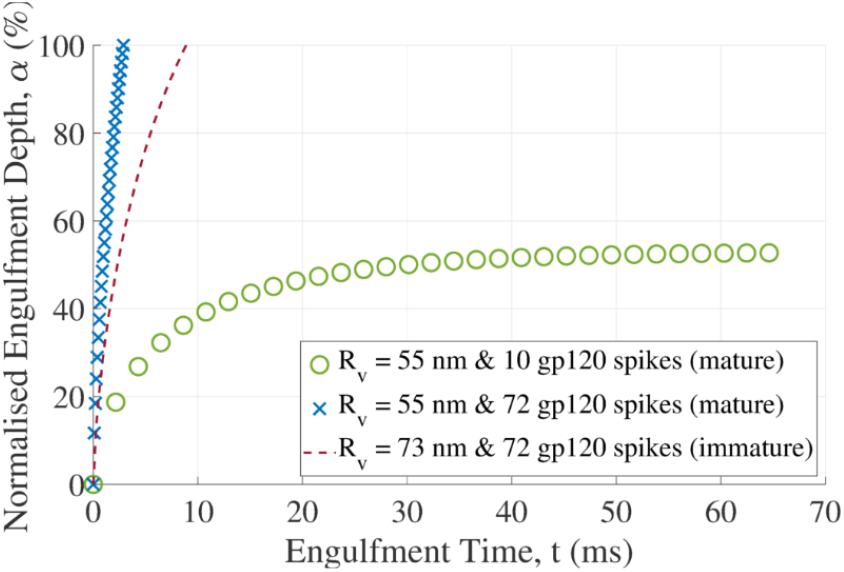
**The normalized engulfment depth of the virion versus engulfment time for HIV particles with different radii and numbers of gp120 spikes. Engulfment is to α = 100% for the immature virion and the mature virion with 72 gp120 spikes ut only to α = 53% for the mature virion with 10 gp120 spikes. A cell radius of *R***_**c**_ **= 4,000 nm was used**

a and b). However, an increase in the magnitude of the normalized total engulfment energy at full engulfment of α = 100% from |-3198.5| to |-3720.9| is predicted for the decrease in virion size (Figure 5 Normalized total engulfment energy versus normalized engulfment depth for HIV particles with different radius and number of gp120 spikes related to virus maturation predicted with (a) the linear-elastic and (b) the viscoelastic model for early-stage engulfment, and with (c) the viscoelastic model for full engulfment. A cell radius of *Rc* = 4,000 nm was used The duration of complete engulfment (α = 100%) is 8.88 ms for the immature virion and 2.88 ms for a mature virion with a smaller size (*Rv* = 55 nm versus *Rv* = 73 nm) but the same number of 72 gp120 spikes as the immature virion (Figure 6). Taking further into account the reduction in the number of gp120 spike proteins (10 versus 72) during maturation, the maximum engulfment is limited to 53% and reached after 64.62 ms.

c).

For the reduction of the number of gp120 spikes from 72 to 10 during maturation, a notable decrease in the normalized energy is predicted even at early-stage engulfment (Figure 5 Normalized total engulfment energy versus normalized engulfment depth for HIV particles with different radius and number of gp120 spikes related to virus maturation predicted with (a) the linear-elastic and (b) the viscoelastic model for early-stage engulfment, and with (c) the viscoelastic model for full engulfment. A cell radius of *Rc* = 4,000 nm was used

The duration of complete engulfment (α = 100%) is 8.88 ms for the immature virion and 2.88 ms for a mature virion with a smaller size (*Rv* = 55 nm versus *Rv* = 73 nm) but the same number of 72 gp120 spikes as the immature virion (Figure 6). Taking further into account the reduction in the number of gp120 spike proteins (10 versus 72) during maturation, the maximum engulfment is limited to 53% and reached after 64.62 ms.

). For the mature virion, a total normalized energy of *W*_Tn_ = |-3198.5| at full engulfment (α = 100%) is predicted for 72 gp120 spikes compared to *W*_Tn_ = |-140.1| for 10 gp120 spikes at a maximum engulfment depth of α = 53% (Figure 5 Normalized total engulfment energy versus normalized engulfment depth for HIV particles with different radius and number of gp120 spikes related to virus maturation predicted with (a) the linear-elastic and (b) the viscoelastic model for early-stage engulfment, and with (c) the viscoelastic model for full engulfment. A cell radius of *Rc* = 4,000 nm was used

The duration of complete engulfment (α = 100%) is 8.88 ms for the immature virion and 2.88 ms for a mature virion with a smaller size (*Rv* = 55 nm versus *Rv* = 73 nm) but the same number of 72 gp120 spikes as the immature virion (Figure 6). Taking further into account the reduction in the number of gp120 spike proteins (10 versus 72) during maturation, the maximum engulfment is limited to 53% and reached after 64.62 ms.

c).

The duration of complete engulfment (α = 100%) is 8.88 ms for the immature virion and 2.88 ms for a mature virion with a smaller size (*R*_v_ = 55 nm versus *R*_v_ = 73 nm) but the same number of 72 gp120 spikes as the immature virion (Figure 6). Taking further into account the reduction in the number of gp120 spike proteins (10 versus 72) during maturation, the maximum engulfment is limited to 53% and reached after 64.62 ms.

## 4. Discussion

A model based on contact mechanics theory was developed to investigate the mechanical virion-cell interactions during the engulfment of the virion and assess the influence of morphological and biophysical parameters of the virion and host cell on the engulfment energy and invagination force. A linear-elastic model previously used for early engulfment with small deformations was compared with a new viscoelastic model for deformations beyond the linear-elastic range of the cellular and viral lipid membranes during full engulfment.

The comparison of the linear-elastic and viscoelastic models indicates that the linear-elastic model can predict the total engulfment energy well for early (α < 10%) and full engulfment (α = 100%). However, the linear-elastic model considerably underestimated the required invagination force for early and full engulfment, demonstrating the advantage of the viscoelastic over the linear-elastic model.

The assessment of cell size and morphology reveals that the variation of the cell radius between lymphocytes and macrophages has minimal impact on the invagination force (Figure 3). This finding aligns with previous reports of the marginal effect of cell radius on the mechanics of the virion-cell interaction (Gefen 2010). However, local morphological surface features of the host cell’s membrane at the nanometre length scale (Gentile et al. 1994) affect the engulfment mechanics substantially. These local surface morphologies with curvatures of the same order of magnitude as those of the virion reduce the invagination force and are potential sites for easier virion entry. This suggests that the surface morphology of host cells may play a role in the infection process and infectivity. The model predicted invagination forces for various cell sizes and morphologies ranging from 11.1 – 28.3 pN. These results are similar to previous experimental quantifications of the interaction strength ranging between 10 – 58 pN (Wiegand et al. 2020; Tsai et al. 2015; Sieben et al. 2012; Pan et al. 2017; Alsteens et al. 2017; Rankl et al. 2008; Sieben et al. 2011).

The required invagination force predicted in the current study was of the same order of magnitude as the adhesion force of 25.45 pN determined experimentally between the gp120 ligand and CD4 cell receptor (Chen et al. 2011).

The decreasing virion size and shedding of gp120 spikes associated with virion maturation affect the engulfment energy. The moderate increase in engulfment energy with the decrease in virion size predicted with the current model agreed with the findings of a previous study (Gefen 2010). A substantial decrease in engulfment energy after shedding gp120 spikes is primarily ascribed to the reduced availability of adhesion energy from gp120. The overall reduction in the engulfment energy during virion maturation suggests a mechanobiological reduction of the virion’s entry ability. While this contradicts previous observations that the entry ability increases as virions mature (Jiang and Aiken 2006; Kol et al. 2007), there may be other determinants of entry ability. One possibility is the conformational change in gp120 during maturation that impacts the functional capabilities of these ligands (Wyma et al. 2004; Murakami et al. 2004). The improved ability of the viral ligands in a mature virion to migrate and cluster in an area of the viral membrane preferential for contact to and engulfment into the host cell can be another mechanism that promotes virion entry (Chojnacki et al. 2017; Chojnacki et al. 2012). The model also does not account for other factors, such as the induction of spontaneous curvatures in the cell membrane due to interactions with virions, which might impact the engulfment process (Nossal and Zimmerberg 2002; Lipowsky and Döbereiner 1998).

Membrane fusion between an HIV particle and the host cell during entry likely occurs at partial engulfment (α < 100%), although it is unknown at which engulfment depth the viral and cell membranes fuse. The final engulfment depth of α = 53% for a mature virion based on the available adhesion energy might be the first estimate of the maximum engulfment depth by which membrane fusion has to occur. As no further engulfment is possible energetically, membrane fusion needs to take place to facilitate the release of viral content into the cell.

The engulfment duration ranged between 2.88 ms for 100% engulfment of a particle with mature virion size before ligand shedding and 64.62 ms for 53% engulfment of the mature virion after shedding of gp120 spikes. The varying engulfment duration may be associated with distinctly different deformation rates of the host cell’s membrane during engulfment and offer an avenue for further studies.

It is widely assumed that engulfment is driven by the adhesion energy gained by cell receptor and virion ligand binding. The model reflects this and is highly sensitive to changes in the receptor density (Figure 5). Reducing the number of ligands from 72 to 10 resulted in an energy decrease from *W*_Tn_ = |-3198.5| to |-140.1| and limited virion engulfment (α = 53%). The model is much less sensitive to changes in virion size. The model is moderately sensitive to large changes in cell size, provided the change is several orders of magnitude (Figure 3).

The current model includes a simplified adhesion energy equation that assumes uniform ligands distribution and a high density of immobile cell receptors. Unlike existing models (Zhdanov 2013, 2015; Bai et al. 2018; Gao et al. 2005; Yi and Gao 2017), the model does not consider receptor diffusion and other entropic or enthalpic contributions. These could be included in future, along with different virion shapes like icosahedral (Katzengold et al. 2016), ellipsoid (Dasgupta et al. 2014), or oblate virions. Other factors, such as the induction of a spontaneous cell membrane curvature due to interaction with the virion, could also be included to enhance further the model’s details (Lipowsky and Döbereiner 1998; Nossal and Zimmerberg 2002).

## Conclusions

This explorative study identified biophysical parameters involved in HIV engulfment as candidates for further research to extend the understanding of the virion entry into host cells. The findings offer the potential for the mechanobiological assessment of the invagination of enveloped viruses towards improving the prevention and treatment of viral infections. The developed simple mathematical model can be extended and advanced by considering, e.g., non-uniform and mobile virion distribution on the virion, cell receptor diffusion, spontaneous membrane curvature induction, and different virion shapes.

## Data Availability

The custom Matlab code files used in this study will be available on the University of Cape Town’s institutional data repository (ZivaHub) under http://doi.org/10.25375/uct.21623142 upon publication.

## Credit Author Contributions

EK: Conceptualisation, Methodology, Investigation, Writing - Original Draft, Writing - Review & Editing

TA: Methodology, Investigation, Writing - Original Draft

PS: Investigation, Writing - Review & Editing

TF: Conceptualisation, Investigation, Project administration, Funding acquisition, Resources, Writing - Review & Editing

## References

Ahadi A, Johansson D, Evilevitch A (2013) Modeling and simulation of the mechanical response from nanoindentation test of DNA-filled viral capsids. J Biol Phys 39:183–199.

Alsteens D, Newton R, Schubert R, Martinez-Martin D, Delguste M, Roska B, Muller DJ (2017) Nanomechanical mapping of first binding steps of a virus to animal cells. Nat Nanotechnol 12:177–183.

Altman YM, Wooford OJ (2014) Export_fig - a toolbox for exporting figures from MATLAB to standard image and document formats. https://github.com/altmany/export_fig. Accessed 29 Nov 2022

Bai F, Wu J, Sun R (2018) An investigation of endocytosis of targeted nanoparticles in a shear flow by a statistical approach. Mathematical Biosciences 295:55–61.

Canham PB (1970) The minimum energy of bending as a possible explanation of the biconcave shape of the human red blood cell. Journal of Theoretical Biology 26:61–81.

Chen Y, Zeng G, Chen SS, Feng Q, Chen ZW (2011) AFM force measurements of the gp120-scd4 and gp120 or cd4 antigen-antibody interactions. Biochem Biophys Res Commun 407:301–306.

Chojnacki J, Staudt T, Glass B, Bingen P, Engelhardt J, Anders M, Schneider J, Muller B, Hell SW, Krausslich HG (2012) Maturation-dependent HIV-1 surface protein redistribution revealed by fluorescence nanoscopy. Science 338:524–528.

Chojnacki J, Waithe D, Carravilla P, Huarte N, Galiani S, Enderlein J, Eggeling C (2017) Envelope glycoprotein mobility on HIV-1 particles depends on the virus maturation state. Nat Commun 8:545.

Choudhary V, Gupta A, Sharma R, Parmar HS (2021) Therapeutically effective covalent spike protein inhibitors in treatment of SARS-CoV-2. Journal of Proteins and Proteomics 12:257–270.

Coomer CA, Carlon-Andres I, Iliopoulou M, Dustin ML, Compeer EB, Compton AA, Padilla-Parra S (2020) Single-cell glycolytic activity regulates membrane tension and HIV-1 fusion. PLOS Pathogens 16:e1008359.

Dasgupta S, Auth T, Gov NS, Satchwell TJ, Hanssen E, Zuccala ES, Riglar DT, Toye AM, Betz T, Baum J, Gompper G (2014) Membrane-wrapping contributions to malaria parasite invasion of the human erythrocyte. Biophysical Journal 107:43–54.

de la Vega M, Marin M, Kondo N, Miyauchi K, Kim Y, Epand RF, Epand RM, Melikyan GB (2011) Inhibition of HIV-1 endocytosis allows lipid mixing at the plasma membrane, but not complete fusion. Retrovirology 8:99.

Deserno M, Bickel T (2003) Wrapping of a spherical colloid by a fluid membrane. Europhysics Letters 62:767.

Dintwa E, Tijskens E, Ramon H (2007) On the accuracy of the Hertz model to describe the normal contact of soft elastic spheres. Granular Matter 10:209–221.

Fregin B, Czerwinski F, Biedenweg D, Girardo S, Gross S, Aurich K, Otto O (2019) High-throughput single-cell rheology in complex samples by dynamic real-time deformability cytometry. Nat Commun 10:415.

Gao H, Shi W, Freund LB (2005) Mechanics of receptor-mediated endocytosis. Proceedings of the National Academy of Sciences 102:9469–9474.

Gefen A (2010) Effects of virus size and cell stiffness on forces, work, and pressures driving membrane invagination in a receptor-mediated endocytosis. J Biomech Eng 132:084501.

Gelderblom HR (1991) Assembly and morphology of HIV: Potential effect of structure on viral function. AIDS 5:617–637.

Gentile M, Adrian T, Scheidler A, Ewald M, Dianzani F, Pauli G, Gelderblom HR (1994) Determination of the size of HIV using adenovirus type-2 as an internal length marker. J Virol Methods 48:43–52.

Gorai B, Sahoo AK, Srivastava A, Dixit NM, Maiti PK (2021) Concerted interactions between multiple gp41 trimers and the target cell lipidome may be required for HIV-1 entry. Journal of Chemical Information and Modeling 61:444–454.

Grief C, Hockley DJ, Fromholc CE, Kitchin PA (1989) The morphology of simian immunodeficiency virus as shown by negative staining electron microscopy. J Gen Virol 70 (Pt 8):2215–2219.

Helfrich W (1973) Elastic properties of lipid bilayers: Theory and possible experiments. znc 28:693–703.

Jiang J, Aiken C (2006) Maturation of the viral core enhances the fusion of HIV-1 particles with primary human t cells and monocyte-derived macrophages. Virology 346:460–468.

Johnson KL (1987) Contact mechanics. Cambridge University Press, Cambridge, UK. doi:10.1017/cbo9781139171731

Katzengold R, Zaharov E, Gefen A (2016) Analytical and computational modeling of early penetration of non-enveloped icosahedral viruses into cells. Technol Health Care 24:483–493.

Kierszenbaum AL, Tres L (2015) Histology and cell biology: An introduction to pathology. 4 edn. Saunders, Philadelphia, PA

Kol N, Shi Y, Tsvitov M, Barlam D, Shneck RZ, Kay MS, Rousso I (2007) A stiffness switch in HIV. Biophys J 92:1777–1783.

Layne SP, Merges MJ, Dembo M, Spouge JL, Conley SR, Moore JP, Raina JL, Renz H, Gelderblom HR, Nara PL (1992) Factors underlying spontaneous inactivation and susceptibility to neutralization of human immunodeficiency virus. Virology 189:695–714.

Leckband D, Israelachvili J (2001) Intermolecular forces in biology. Q Rev Biophys 34:105–267.

Li L, Zhang Y, Wang J (2018) A calibration model on dynamic measurements of nanoparticles endocytosis. EPL (Europhysics Letters) 124.

Lipowsky R, Döbereiner HG (1998) Vesicles in contact with nanoparticles and colloids. Europhysics Letters (EPL) 43:219–225.

Micoulet A, Spatz JP, Ott A (2005) Mechanical response analysis and power generation by single-cell stretching. Chemphyschem 6:663–670.

Miyauchi K, Kim Y, Latinovic O, Morozov V, Melikyan GB (2009) HIV enters cells via endocytosis and dynamin-dependent fusion with endosomes. Cell 137:433–444.

Murakami T, Ablan S, Freed EO, Tanaka Y (2004) Regulation of human immunodeficiency virus type 1 Env-mediated membrane fusion by viral protease activity. Journal of Virology 78:1026–1031.

Nossal R, Zimmerberg J (2002) Endocytosis: Curvature to the enth degree. Curr Biol 12:R770–772.

Pan Y, Zhang F, Zhang L, Liu S, Cai M, Shan Y, Wang X, Wang H, Wang H (2017) The process of wrapping virus revealed by a force tracing technique and simulations. Adv Sci (Weinh) 4:1600489.

Pang HB, Hevroni L, Kol N, Eckert DM, Tsvitov M, Kay MS, Rousso I (2013) Virion stiffness regulates immature HIV-1 entry. Retrovirology 10:4.

Rankl C, Kienberger F, Wildling L, Wruss J, Gruber HJ, Blaas D, Hinterdorfer P (2008) Multiple receptors involved in human rhinovirus attachment to live cells. Proc Natl Acad Sci U S A 105:17778–17783.

Sieben C, Kappel C, Wozniak A, Zhu R, Rankl C, Hinterdorfer P, Grubmüller H, Herrmann A (2011) Influenza virus adhesion to living cells measured by single virus force spectroscopy (svfs) and force probe md simulation. Biophysical Journal 100:22a–23a.

Sieben C, Kappel C, Zhu R, Wozniak A, Rankl C, Hinterdorfer P, Grubmuller H, Herrmann A (2012) Influenza virus binds its host cell using multiple dynamic interactions. Proc Natl Acad Sci U S A 109:13626–13631.

Sun SX, Wirtz D (2006) Mechanics of enveloped virus entry into host cells. Biophys J 90: L10–12.

Tsai B-Y, Chen J-Y, Chiou A, Ping Y-H (2015) Using optical tweezers to quantify the interaction force of dengue virus with host cellular receptors. Microscopy and Microanalysis 21:219–220.

Wiegand T, Fratini M, Frey F, Yserentant K, Liu Y, Weber E, Galior K, Ohmes J, Braun F, Herten D-P, Boulant S, Schwarz US, Salaita K, Cavalcanti-Adam EA, Spatz JP (2020) Forces during cellular uptake of viruses and nanoparticles at the ventral side. Nature Communications 11:32.

Wilen CB, Tilton JC, Doms RW (2012) HIV: Cell binding and entry. Cold Spring Harbor Perspectives in Medicine 2.

Wyma DJ, Jiang J, Shi J, Zhou J, Lineberger JE, Miller MD, Aiken C (2004) Coupling of human immunodeficiency virus type 1 fusion to virion maturation: A novel role of the gp41 cytoplasmic tail. Journal of Virology 78:3429–3435.

Yi X, Gao H (2017) Kinetics of receptor-mediated endocytosis of elastic nanoparticles. Nanoscale 9:454–463.

Zaitseva E, Zaitsev E, Melikov K, Arakelyan A, Marin M, Villasmil R, Margolis LB, Melikyan GB, Chernomordik LV (2017) Fusion stage of HIV-1 entry depends on virus-induced cell surface exposure of phosphatidylserine. Cell Host & Microbe 22:99–110.e117.

Zhdanov VP (2013) Physical aspects of the initial phase of endocytosis. Physical Review E 88:064701.

Zhdanov VP (2015) Kinetics of virus entry by endocytosis. Physical Review E 91:042715.

